# E-SNPs&GO: Embedding of protein sequence and function improves the annotation of human pathogenic variants

**DOI:** 10.1101/2022.05.10.491314

**Authors:** Matteo Manfredi, Castrense Savojardo, Pier Luigi Martelli, Rita Casadio

## Abstract

**Motivation:** The advent of massive DNA sequencing technologies is producing a huge number of human single-nucleotide polymorphisms occurring in protein-coding regions and possibly changing protein sequences. Discriminating harmful protein variations from neutral ones is one of the crucial challenges in precision medicine. Computational tools based on artificial intelligence provide models for protein sequence encoding, bypassing database searches for evolutionary information. We leverage the new encoding schemes for an efficient annotation of protein variants.

**Results:** E-SNPs&GO is a novel method that, given an input protein sequence and a single residue variation, can predict whether the variation is related to diseases or not. The proposed method, for the first time, adopts an input encoding completely based on protein language models and embedding techniques, specifically devised to encode protein sequences and GO functional annotations. We trained our model on a newly generated dataset of 65,888 human protein single residue variants derived from public resources. When tested on a blind set comprising 6,541 variants, our method outperforms recent approaches released in literature for the same task, reaching a MCC score of 0.71. We propose E-SNPs&GO as a suitable, efficient and accurate large-scale annotator of protein variant datasets.

## 1 Introduction

Single-nucleotide polymorphisms (SNPs) are major drivers of human evolution. In many cases, these variations can be directly associated with the onset of genetic diseases. Specifically, SNPs occurring in protein-coding regions often lead to observable changes in the protein residue sequence. Single-Residue Variations (SRVs) may have an impact at different levels, hampering protein structure, function, stability, localization and interaction with other proteins and/or nucleotides, hence setting the basis for the onset of pathologic conditions (Vihinen, 2021; Lappalainen and McArthur, 2021 and references therein).

Public databases such as HUMSAVAR (The UniProt Consortium, 2021) and ClinVar (Landrum *et al*., 2018) store a compendium of known SRVs and provide, when available, information about the variant clinical significance. However, clear associations to diseases are still unknown for many SRVs, which substantially remain of Uncertain Significance (US). Therefore, SRV annotation is an issue, and effective computational tools are needed to provide large-scale annotation of uncharacterized human variation data.

In the past years, several computational approaches have been implemented, with the aim of annotating whether a protein variation is or not disease associated (Ng and Henikoff, 2001; Li *et al*., 2009; Calabrese *et al*., 2009; Adzhubei *et al*., 2010; Schwarz *et al*., 2010; Choi *et al*., 2012; Carter *et al*., 2013; Niroula *et al*., 2015; Jagadeesh *et al*., 2016; Raimondi *et al*., 2017; Pejaver *et al*., 2019; Yang *et al*., 2022). Methods like SIFT (Ng and Henikoff, 2001) or PROVEAN (Choi *et al*., 2012) are based on the conservation analysis in multiple sequence alignments. More complex approaches stand on different types of machine-learning frameworks. These include neural networks (Pejaver *et al*., 2019), random forests (Li *et al*., 2009; Carter *et al*., 2013; Niroula *et al*., 2015; Raimondi *et al*., 2017; Yang *et al*., 2022), gradient tree boosting (Jagadeesh *et al*., 2016), support vector machines (Calabrese *et al*., 2009) and naive Bayes classifiers (Adzhubei *et al*., 2010; Schwarz *et al*., 2010). Each method is trained/tested on different datasets of SRVs, either extracted directly from public resources like HUMSAVAR (The UniProt Consortium, 2021) and/or ClinVar (Landrum *et al*., 2018), or taking advantage of pre-compiled datasets of variations, like VariBench (Nair and Vihinen, 2013). Different types of descriptors extract salient features of the protein sequence and/or the local sequence context surrounding the variant position, including physicochemical properties, sequence profiles, conservation scores, predicted structural motifs and functional annotations. SNPs&GO (Calabrese *et al*., 2009) firstly recognized the importance of functional annotations for the prediction of variant pathogenicity and introduced the LGO feature, a score of association between Gene Ontology (GO) annotations and the variant pathogenicity. The incorporation of the LGO feature significantly improved the prediction performance of SNPs&GO (Calabrese *et al*., 2009).

Recent developments in the field of deep learning focus on the definition of new ways of representing protein sequences. Large-scale protein language models are inspired and derived from the Natural Language Processing field (Ofer *et al*, 2021). They learn numerical vector representations of protein sequences, containing important features that are reflected in the evolutionary conservation and in the sequence syntax (Bepler and Berger, 2021). These numerical vectors are then adopted to encode protein sequence and/or individual residues in place of canonical, hand-crafted features such as physico-chemical properties or evolutionary information. These distributed protein representations emerge from the application of generative learning models trained on large databases of sequence data (Bepler and Berger, 2021; Ofer et al, 2021; Ibtehaz and Kihara, 2021)

Successful protein language models are routinely trained on databases composed of hundreds of millions of unique sequences with hundreds of billions of residues. Training is computationally demanding, routinely requiring weeks or months of computations on high-performance Graphical Processing Units (GPUs) (Rives *et al*., 2021; Elnaggar *et al*., 2021; Vig *et al*., 2021). However, the advantage is that most of the computational cost is concentrated on the training phase, and once the model/s is/are trained they can be adopted to embed new sequences with limited resources both in terms of time, memory and computational power.

Embeddings obtained with language models have been recently employed for many different applications with great success (Ibtehaz and Kihara, 2021), including the prediction of protein function and localization (Stärk *et al*., 2021; Littmann *et al*., 2021; Teufel *et al*. 2022), of protein contact maps (Singh *et al*., 2022) and binding sites (Mahbub and Bayzid, 2022).

Several pre-trained language models currently exist in the literature (Asgari and Mofrad, 2015; Strodthoff *et al*., 2020; Alley *et al*., 2019; Heinzinger *et al*., 2019; Rives *et al*., 2021; Elnaggar *et al*., 2021), mainly differing in the specific generative architectures (autoregressive, bidirectional, masked; Bepler and Berger, 2021) and datasets adopted for training.

Not limited to the encoding of protein sequence data, embedding techniques are also applied to model the relationships existing within more complex structures such as graphs, networks, or biological ontologies (Grover and Leskovec, 2016; Perozzi *et al*., 2014; Zhong *et al*., 2020; Kandathil *et al*., 2022; Edera *et al*., 2022).

In this paper we attempt to fully exploit the power of language models and embeddings for the prediction of variant pathogenicity from the human protein sequence. On the methodological side, two major contributions can be highlighted. Firstly, we adopt two different and complementary embedding procedures, ProtTrans T5 (Elnaggar *et al*., 2021) and ESM-1v (Meier *et al*., 2021), to directly encode an input variation without introducing any hand-crafted feature as previously done. Secondly, leveraging the idea introduced in SNPs&GO (Calabrese *et al*., 2009), we explore a new way of encoding functional annotations by adopting a model called Anc2Vec (Edera *et al*., 2022), specifically designed for the embedding of Gene Ontology (GO) terms.

We trained a Support Vector Machine (SVM) using the above input encoding on a newly generated dataset of 65,888 human disease-related and benign variations obtained from the rational merging of data deposited in two databases, HUMSAVAR (The UniProt Consortium, 2021) and ClinVar (Landrum *et al*., 2018). The method is tested on an independent, non-redundant blind set comprising 6,541 variations adopting stringent homology reduction and evaluation procedures. Results obtained in a comparative benchmark and including one of the most recent and effective methods (Pejaver *et al*., 2019), demonstrate that our model performs at the level or even better than the state-of-the-art, reaching a MCC of 0.71. Based on an input encoding derived solely from embedding models, our method is fast: this makes it suitable for large-scale annotation of human pathogenic variants.

## 2 Methods

### 2.1 Dataset

We obtained the dataset of Single Residue Variations (SRVs) by merging information extracted from two resources: HUMSAVAR (accessed on Aug 4th, 2021), listing all missense variants annotated in human UniProt/Swiss-Prot (The UniProt Consortium, 2021) entries, and ClinVar (accessed on March 29th, 2021), the NCBI resource of relationships among human variations and disease phenotypes (Landrum *et al*., 2018).

Both databases classify the effect of SRVs into different classes: Pathogenic or Likely Pathogenic (P/LP), Benign or Likely Benign (B/LB), and of Uncertain Significance (US). We retained only P/LP SRVs clearly associated with the diseases catalogued in OMIM (Amberger *et al*., 2019) or in MONDO (Shefchek *et al*., 2020). We collected also all the B/LB variations and excluded SRVs labelled as US, somatic, or with contrasting annotations of the effect.

Overall, the dataset consists of 11,565 protein sequences endowed with 72,429 SRVs, including 30,481 P/LP SRVs in 2873 proteins and 41,948 B/LB SRVs in 11,166 proteins (Table 1, last row).

**Table 1.** The dataset of Single Residue Variations (SRVs) adopted in this study

| Dataset | # of pathogenic SRVs | # of neutral SRVs | # of proteins |
| --- | --- | --- | --- |
| Training set | 27,734 | 38,154 | 10,533 |
| Blind test set | 2,747 | 3,794 | 1,032 |
| Total | 30,481 | 41,948 | 11,565 |

For all proteins in the dataset, we extracted Gene Ontology (GO) annotations from the corresponding entry in UniProt. Overall, our dataset is annotated with 16,092 GO terms, including 10,817 BP (Biological Process), 3,766 MF (Molecular Function) and 1,509 CC (Cellular Component).

#### 2.1.1 Cross-validation and generation of the blind test set

To avoid biases between training and testing sets, we adopted a stringent clustering procedure to generate cross-validation sets. Firstly, we clustered protein sequences with the MMseqs2 program (Steinegger and Söding, 2017), by constraining a minimum sequence identity of 25% over a pairwise alignment coverage of at least 40%. We used a connected component clustering strategy so that if two proteins are clustered with a third one, they all end up in the same set. We then split our variants into 11 equally distributed subsets, making sure that all variations, occurring in proteins that clustered together, are part of the same subset (Supplementary Table 1). Ten out of 11 subsets are used in a 10-fold cross-validation procedure for optimizing the input encoding and for fixing the model hyperparameters. The remaining subset is the blind test set for assessing the generalization performance of our approach and for benchmarking it with other popular methods available in literature. In this way, we limit sequence redundancy between training and testing sets, enabling a fair evaluation of the results. It is worth noticing that the blind test can share similarity with proteins included in the training sets of the other benchmarked methods.

### 2.2 General overview of the approach

Figure 1 depicts the architecture of E-SNPs&GO, including an input encoding, a predictor and an output. The input consists of a human protein sequence and a single residue variation occurring at a specific position along the sequence. In the input encoding phase, the sequence and its variant are embedded with two different procedures, ESM-1v (Meier *et al*., 2021) and ProtTrans T5 (Elnaggar et al., 2021). In order to embed the functional protein annotation of the wildtype protein, we adopt Anc2Vec (Edera *et al*., 2022).

**Figure 1.**
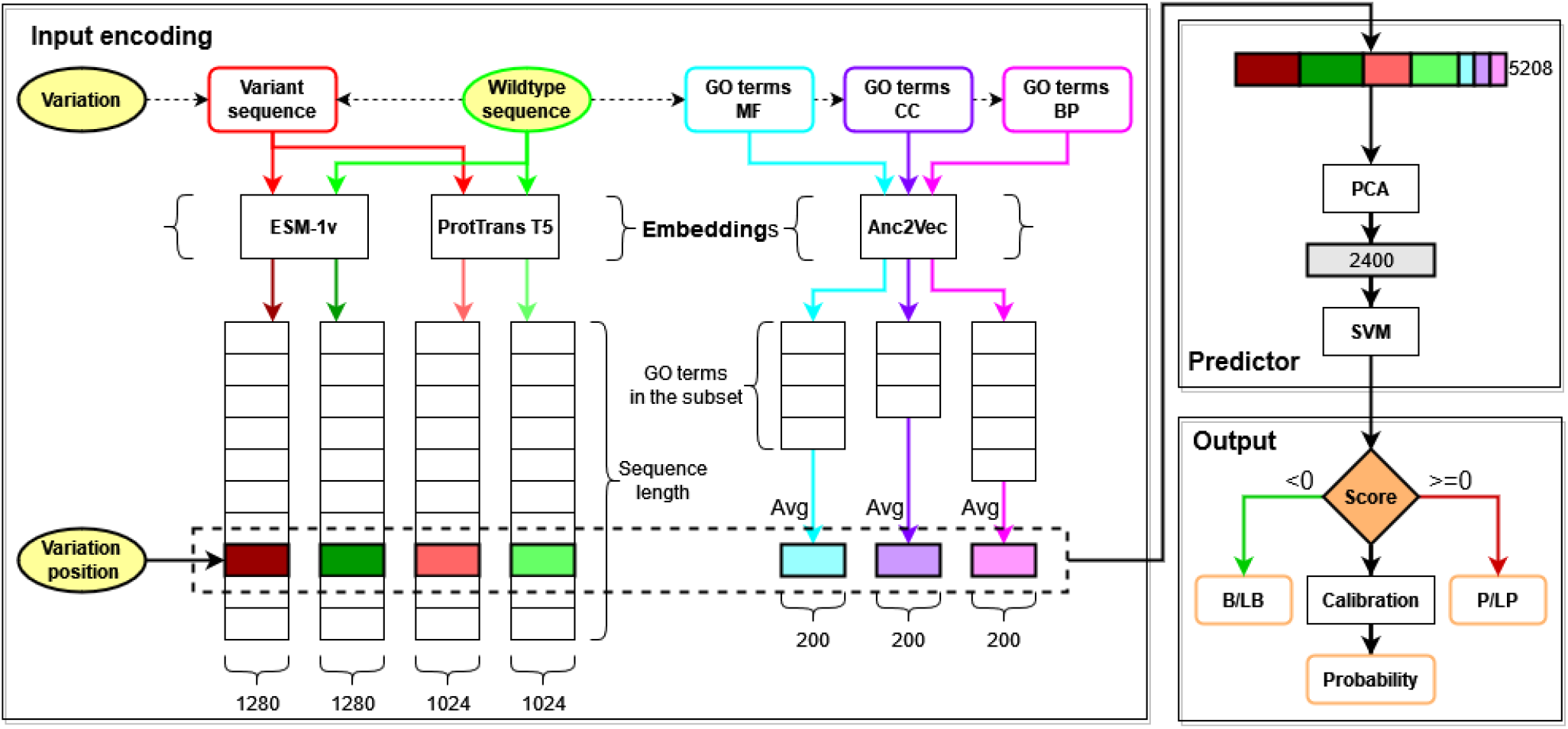
General overview of the architecture of E-SNPs&GO. Inputs (wildtype sequence, variation and variation position) are in yellow. The architecture includes an Input encoding, a Predictor and an Output. During the Input encoding, three embedding models are adopted to generate vector representations. The wildtype sequence (green) and the variant sequence (red) are modelled with ESM-1v (Meier et al., 2021) and ProtTrans T5 (Elnaggar et al. 2021). The GO functional annotations (blue MF (Molecular Function), purple CC (Cellular Component), pink BP (Biological Process)) are modelled with Anc2Vec (Edera et al., 2022). The vectors within the dashed box (marked with different colours), representing the variation position and the averaged (Avg) GO terms of the wildtype sequence, are then concatenated together to obtain a final representation consisting of 1280*2 + 1024*2 + 200*3 = 5208 features. This vector is fed to the Predictor, which includes a Principal Component Analysis (PCA) to reduce the input dimensionality (from 5208 to 2400) and a Support Vector Machine (SVM) performing as Output a binary classification into Benign/Likely Benign (B/LB, negative class, Score < 0) or Pathogenic/Likely Pathogenic (P/LP, positive class, Score >= 0). We apply an Isotonic Regression (Calibration) to obtain a calibrated probability.

In the predictor, the vector representation generated in the input encoding is then processed using a Principal Component Analysis (PCA), which reduces the dimensionality of the input. The output feeds a Support Vector Machine (SVM) classifier performing the final labelling as Pathogenic (P/LP) or Benign (B/LB). A given input variant is predicted as pathogenic when the SVM output score is greater than or equal to zero, benign otherwise. A final calibration step allows to convert scores into probabilities for a variant to be pathogenic. Details of the methods included in E-SNPs&GO, are listed in the following sections.

### 2.3 Input encoding: embeddings of protein sequence, its variant and GO terms

#### 2.3.1 Transformers for embedding of protein sequences and their variants

Several prominent language models and corresponding embedding generation schemes in Natural Language Processing (NLP) are available, and some of these have been adapted to protein sequences to perform specific prediction tasks (Ibtehaz and Kihara, 2021). Large-scale Protein Language Models (PLMs) aim at learning a numerical vector representation that allows reconstructing the input sequence. Briefly, PLMs are trainable generative models aiming at reconstructing masked residues of a protein sequence starting from the context. They consist of two modules: i) the feature encoding module (encoder) provides a non-linear representation of the input (context) and ii) the generative module (decoder) reconstructs the masked portion. Both modules are trained together on a large set (routinely in the range of hundreds of millions) of real protein sequences. The learned representation captures important features of the proteins, including physicochemical, structural, functional, and evolutionary features (Bepler and Berger, 2021; Ofer *et al*., 2021). Depending on the problem at hand, after the training the generative part of the model (decoder) can be neglected, keeping only the encoding function that allows to compute the embedding of new sequences. By transfer learning, the embedded schemes can be provided as input to machine- and deep-learning architectures for a predictive task. Among PLMs, transformer-based masked models (Vaswani *et al*., 2017), aim to solve the problem of efficiently capturing long-distance interactions in the sequence. Transformers are architectures that include a self-attention mechanism to extract the context information from the whole sequence (Vaswani et al., 2017).

In this paper we adopt two different transformers for protein embedding schemes, ESM-1v (Meier *et al*., 2021) and ProtTransT5 (Elnaggar *et al*., 2021). Both embedding methods have been developed and tested on protein variant related problems (Meier *et al*., 2021; Marquet *et al*., 2021). The major difference stands in the volume of the sequence datasets used for generating the embedding schemes and in the adoption of different training procedures. ESM-1v was trained on a single run using a dataset of 98 million unique sequences extracted from UniRef90 (Suzek *et al*., 2015). ProtTrans T5 (version XL U50) was trained using a two-step procedure: in a first pass, training was performed using the large BFD database (Steinegger *et al*., 2019; Steinegger and Söding, 2018), comprising the whole UniProt as well as protein sequences translated from multiple metagenomic sequencing projects, and consisting of about 2.1 billion unique sequences. In the second pass, a fine-tuning of the model was obtained using a smaller database derived from UniRef50 (Suzek *et al*., 2015) and including 45 million unique sequences.

#### 2.3.2 Embedding of biological ontologies

The concept of embedding can be generalised to any kind of data with different underlying structures such as graphs or networks (Grover and Leskovec, 2016; Perozzi *et al*., 2014). In particular, several embedding models, have been defined to provide a numerical representation of nodes in ontologies (Zhong *et al*., 2020; Chen *et al*., 2021). Here we adopt Anc2Vec (Edera *et al*., 2022), a method that learns a vector representation for GO terms, by preserving ancestors relationships.

Because the embedding is not context-dependent, we precompute the vector representation for each possible GO term.

### 2.4 Predictor input

For encoding variations, we firstly perform a full-sequence generation of embeddings using both the ESM-1v (Meier *et al*., 2021) and the ProtTrans T5 XL U50 (Elnaggar *et al*., 2021) models. Given a protein sequence with *L* residues, this provides protein encodings of dimensions *L*x1280 and *L*x1024, respectively. Sequence embeddings are carried out independently on both the wild-type and the variant sequence.

For a variation in position *i* in a protein sequence we compute a vector of 4608 features, including:

- 1280 features corresponding to ESM-1v embedding in position *i* of the variated sequence.
- 1280 features corresponding to ESM-1v embedding in position *i* of the wild-type sequence.
- 1024 features corresponding to ProtTrans T5 (version XL U50) embedding in position *i* of the variated sequence.
- 1024 features corresponding to ProtTransT5 (version XL U50) embedding in position *i* of the wild-type sequence.

The ESM-1v embedding model constrains the maximal protein length (*L*) to 1024 residues. For this reason, variations occurring on longer sequences were encoded using a 201 long sequence window centered on the variant position.

After this step, we extract all the GO terms annotated in the UniProt entry of the wildtype protein carrying the variation. Potential term redundancy is removed by retaining only leaf terms. Terms from the three different GO sub-ontologies (Molecular Function, Cellular Component, Biological Process) are processed independently. Each annotated GO term is then embedded as a vector of 200 features using the Anc2Vec model (Edera *et al*., 2022). To obtain a single vector representation independent of the number of terms of a given protein, we average all the vector encodings (Figure 1). Three final average vectors, one for each GO sub-ontology, are concatenated obtaining a protein function encoding of 600 components.

The final variation encoding comprises 5,208 features, obtained by merging the local positional embedding (4,608 features from ESM-1V + ProtTrans T5 XL U50) described above and the Anc2Vec functional encoding (600 features). Eventually, we encode the different embeddings separately (see Results, Table 2).

**Table 2.** Performance of different embedding schemes

| Input encoding | Q<sub>2</sub><br>(%) | Precision<br>(%) | Recall<br>(%) | F1-score<br>(%) | ROC-AUC<br>(%) | MCC |
| --- | --- | --- | --- | --- | --- | --- |
| ESM-1v | 82.4 (±1.5) | 80.4 (±2.6) | 77.0 (±2.8) | 78.6 (±1.9) | 81.6 (±1.5) | 0.637 (±0.031) |
| ESM-1+GO | 83.3 (±1.4) | 81.7 (±2.5) | 78.1 (±2.7) | 79.8 (±1.8) | 82.6 (±1.4) | 0.657 (±0.029) |
| ProtTrans T5 | 83.0 (±1.3) | 79.8 (±1.9) | 80.0 (±2.8) | 79.9 (±1.7) | 82.6 (±1.4) | 0.653 (±0.027) |
| ProtTrans T5+GO | 83.7 (±1.1) | 81.8 (±1.9) | 79.2 (±2.5) | 80.5 (±1.5) | 83.1 (±1.3) | 0.674 (±0.024) |
| ESM-1v+ProtTrans T5 | 83.6 (±1.4) | 81.8 (±2.3) | 78.6 (±2.9) | 80.1 (±1.8) | 82.9 (±1.5) | 0.662 (±0.029) |
| ESM-1v+ProtTrans T5+GO | 84.5 (±1.3) | 82.8 (±2.4) | 79.7 (±2.4) | 81.2 (±1.6) | 83.8 (±1.3) | 0.680 (±0.026) |

ESM-1v (2x1280 = 2560 features);
ESM-1v + GO (2x1280 + 3x200 = 3160 features);
ProtTrans T5 (2x1024 = 2048 features);
ProtTrans T5 + GO (2x1024 + 3x200 = 2648 features);
ESM-1v + ProtTrans T5 (2x1280 + 2x1024 = 4608 features);
ESM-1v + ProtTrans T5 + GO (2x1280 + 2x1024 + 3x200 = 5208 features).
For mean values computed over the 10 cross-validation sets we report standard deviations within brackets.
Scoring indexes are defined in section 2.6

### 2.5 Model selection and implementation

The predictor includes two cascading components (Figure 1): a Principal Component Analysis (PCA) for reducing the dimensionality of the input features and a binary Support Vector Machine (SVM) with a Radial Basis Function (RBF) kernel, which performs the variant classification into pathogenic or not. We optimized the hyperparameters of both methods (such as the number of components of PCA, the SVM cost parameter C and the gamma coefficient of the RBF kernel) with a grid search procedure. A complete list of hyperparameters tested and their optimal values are available in Supplementary Table 2.

It is worth clarifying that, during both cross-validation and blind testing, the execution of the PCA step is always computed on the training set and then applied for projecting vectors of the testing set in the reduced space.

All methods are implemented in Python3 using the scikit-learn library (Pedregosa *et al*., 2011). ESM-1v and ProtTrans T5 embeddings are computed with the bio-embeddings package (Dallago *et al*., 2021).

The complete machine-learning workflow is compliant with the DOME recommendation checklist (Walsh et al., 2021), as reported in Supplementary Table 3.

### 2.6 Scoring indexes

We assess the performance with the following scores. Pathogenic/Likely Pathogenic variations are assumed to be the positive class, while Benign/Likely Benign variations are the negative class. In what follows, *TP, TN, FP* and *FN* are true positive, true negative, false positive and false negative predictions, respectively.

We compute the following scoring measures:

- Accuracy (Q2):

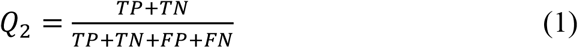
- Precision:

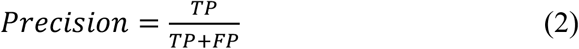
- Recall:

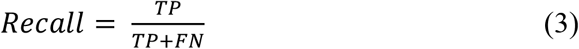
- F1-score, the harmonic mean of precision and recall:

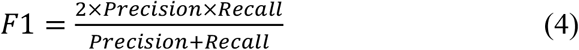
- Area Under the Receiver Operating Characteristic Curve (ROC-AUC).
- Matthews Correlation Coefficient (MCC):

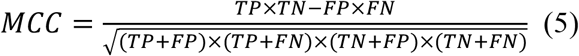

### 2.7 Output calibration

The SVM adopted for classification computes a decision function that represents the distance of the point mapping the input from the discrimination boundary. We use this value to estimate the reliability of the prediction, in terms of the probability of the input variation to be pathogenic (Figure 1)

In a perfectly calibrated method, when a set of predictions scored with probability P is tested on real data, we expect that the fraction of true positives is exactly P. In this work, we adopt a procedure previously described (Benevenuta et al., 2020) to obtain a calibrated probability that we provide in output alongside the predicted class. In particular, we fit an Isotonic Regression (Niculescu-Mizil and Caruana, 2005) in cross-validation and we use it to obtain a probability score on the blind test.

## 3 Results

### 3.1 Assessing the contribution of different input encodings

To select the optimal input encoding, we performed different experiments to test various combinations of input features. To this aim, we trained in cross-validation several independent SVM+PCA models using different input features and using MCC to score and select the optimal model.

GO terms provide global protein information. Their embedding does not consider the specific variant position. If the prediction is run considering only averaged embedded GO terms vector (Figure 1), the predictor performance is very low (MCC=0.27, data not shown). Different input encodings, corresponding to different predictors, perform differently (Table 2). The inclusion of GO embeddings in the final input is always beneficial, improving MCC by 2 or 3 percentage points in all cases (compare ESM-1v, ProtTrans T5 and ESM-1v+ ProtTrans T5 with or without GO, respectively in Table 2). Considering the two protein sequence embeddings, ProtTrans T5 outperforms ESM-1v both with and without the additional GO information. Most notably, the model trained on data from ProtTrans T5 alone is the most balanced, reaching equal precision and recall. Finally, the concatenation of both sequence encodings and the GO embedding provides the best performance (MCC=0.68), leading to an increase in precision without a corresponding decrease in recall.

Based on these results, we select the model trained with ESM-1v+ProtTrans T5+GO as the optimal one.

### 3.2 Benchmark on the blind test set

We test our method adopting both a 10-fold cross-validation procedure and an independent blind test set constructed to be non-redundant with respect to the training dataset (see Section 2.1). Table 3 lists the results. E-SNPs&GO obtains similar results in cross-validation and blind test, making it very robust to generalisation. Concerning individual indexes, our method seems to be slightly more precise than sensitive (compare Precision and Recall).

**Table 3.** Benchmark of our and other top scoring methods available in literature

| Input encoding |  | Q<sub>2</sub> | Precision | Recall | F1-score | ROC-AUC | MCC |
| --- | --- | --- | --- | --- | --- | --- | --- |
|  |  | (%) | (%) | (%) | (%) | (%) |  |
| E-SNPs&GO | Cross-validation | 84.5 | 82.8 | 79.7 | 81.2 | 83.8 | 0.68 |
| E-SNPs&GO | Blind test set | 86.0 | 85.3 | 80.4 | 82.8 | 85.2 | 0.71 |
| SNPs&GO | Blind test set | 79.8 | 84.8 | 63.2 | 72.4 | 77.5 | 0.58 |
| MutPred2.0 | Blind test set | 85.4 | 79.8 | 87.1 | 83.3 | 85.6 | 0.70 |
| PROVEAN | Blind test set | 77.9 | 70.6 | 81.1 | 75.5 | 78.4 | 0.56 |
| PolyPhen-2 | Blind test set | 74.6 | 64.0 | 90.6 | 75.0 | 76.8 | 0.54 |
| SIFT | Blind test set | 74.8 | 65.2 | 85.7 | 74.1 | 76.4 | 0.52 |

Cross-validation is 10-fold and performed on our training set comprising 65,888 variations
For mean values computed over the 10 cross-validation sets we report standard deviations within brackets.
Scoring indexes are defined in section 2.6

Table 3 includes also a comparative benchmark of our method with other state-of-the-art tools, including our SNPs&GO (Calabrese *et al*., 2009), SIFT (Ng and Henikoff, 2001), PolyPhen-2 (Adzhubei et al., 2010), PROVEAN (Choi *et al*., 2012) and MutPred2 (Pejaver *et al*., 2019), one of the most recent and best-performing approaches in the field. Methods are scored adopting our blind test set (Section 2.1), ensuring a fair evaluation of the performance of our method. However, this does not completely exclude the presence of biases in the evaluation of the other tools, since variations included in our blind test may be present in the respective training sets, leading to potential overestimation of their performance.

From Table 3, it is evident that in this benchmark our method is performing at the state-of-the-art. Among tested approaches, PROVEAN, PolyPhen-2 and SIFT, reporting MCCs of 0.56, 0.54 and 0.52 respectively, are scoring lower than our previous SNPs&GO (that achieves an MCC of 0.58). Our E-SNPs&GO and MutPred2, score with significantly higher MCC values of 0.71 and 0.70, respectively. Noticeably the embedding procedure seems to grasp all the properties extracted by an ensemble of different predictors of functional, structural and physicochemical properties, such as the one used by MutPred2 (including 53 tools). Looking at individual scoring measures, MutPred2 appears more sensitive while our method reports a higher precision.

A detailed ablation study performed to evaluate the effect of the GO terms on the prediction scores (Supplementary Table 4), indicates that the most relevant GO subontology is the Molecular Function.

### 3.3 Output probability and reliability index

We associate to each predicted classification (Pathogenic/Likely Pathogenic vs Benign/Likely Benign) a probability score (P) that evaluates the prediction confidence. We adopt a calibration procedure to compute the probability (Benevenuta *et al*., 2020) and this can directly estimate the True Positive Rate (TPR) of a set of predictions with the same score P. For a perfectly calibrated method, P=TPR. Figure 2 plots the calibration curves (TPR vs P) for our and other methods which provide a probability output when tested on the blind test set.

**Figure 2.**
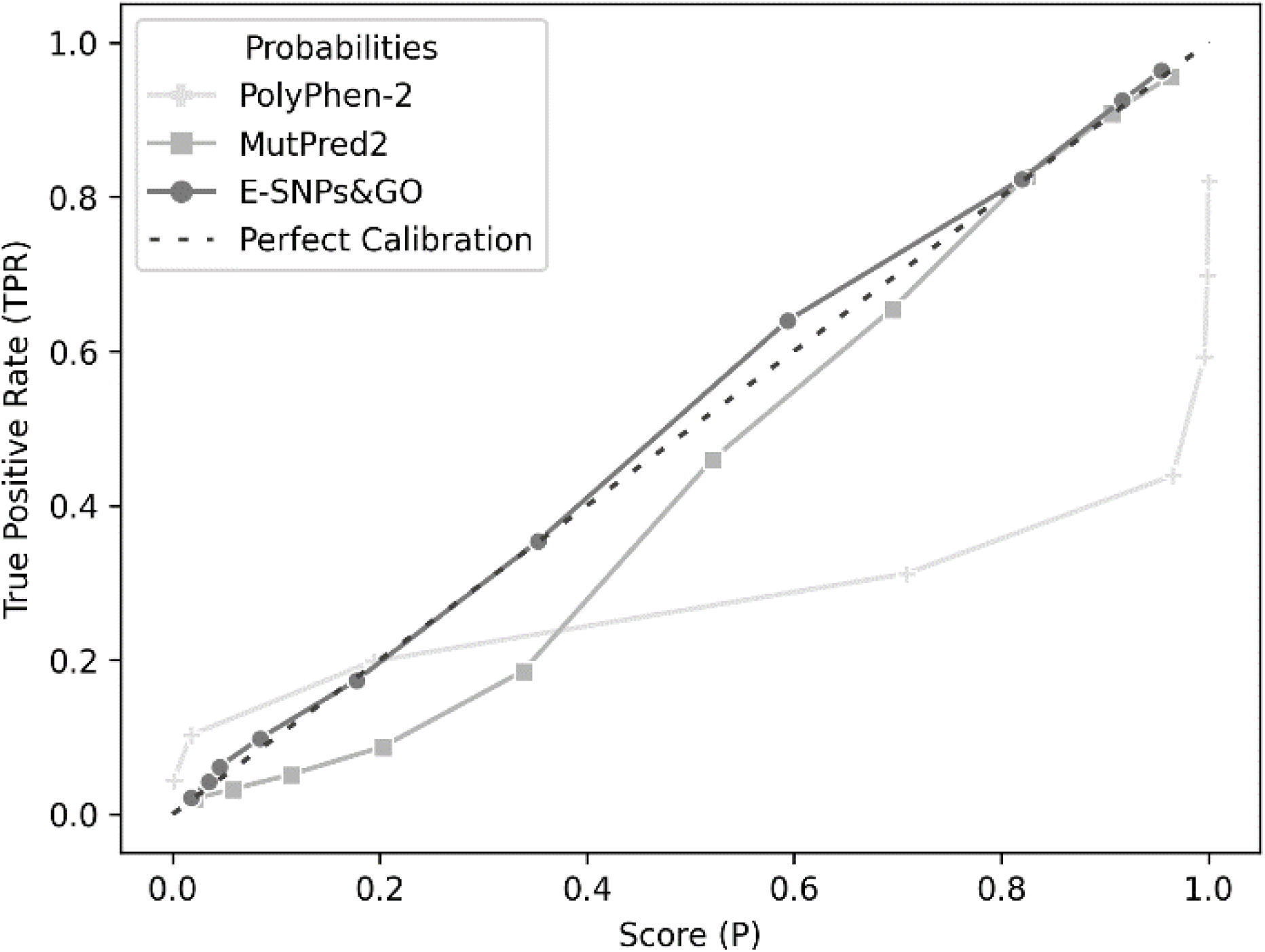
Calibration curves of different methods giving output probabilities on our blind test set. For E-SNPs&GO, the calibration procedure is based on the predictions computed on the training set, and applied to the blind set, preventing information leak.

The dark grey line shows that E-SNPs&GO results are very close to the diagonal line, indicating perfect calibration. MutPred2 is also fairly calibrated, while the curve of PolyPhen-2 is very far from the ideal calibration.

The probability score (P) gives a Reliability Index from 0 (random prediction) to 1 (certain prediction) using the formula:

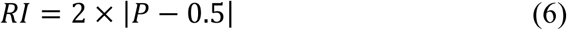

