## Supplementary Table for "E-SNPs&GO: Embedding of protein sequence and function improves the annotation of human pathogenic variants"

**Supplementary Table 1**. Composition of the 10 cross-validation (CV) subsets of E-SNPs&GO in terms of number of proteins and of Pathogenic/Likely Pathogenic (P/LP) and Benign/Likely Benign (B/LB) Single Residue Variations (SRV). Data available at […]

| **CV subset** | **# of P/LP SRVs** | **# of B/LB SRVs** | **# of proteins** |
| --- | --- | --- | --- |
| 1 | 2783 | 3811 | 1057 |
| 2 | 2768 | 3805 | 1044 |
| 3 | 2830 | 3845 | 1127 |
| 4 | 2895 | 3880 | 1103 |
| 5 | 2750 | 3811 | 1043 |
| 6 | 2747 | 3794 | 1032 |
| 7 | 2740 | 3806 | 1035 |
| 8 | 2742 | 3799 | 1030 |
| 9 | 2736 | 3805 | 1030 |
| 10 | 2743 | 3798 | 1032 |

**Supplementary Table 2.** Model hyperparameters optimized with grid search.

| Model | Parameter name | Tested values | Optimal value |
| --- | --- | --- | --- |
| PCA | Number of components | 100, 300, 900, 1200, 1800, 2400, 3000, 5208 | 2400 |
| SVM | C (cost parameter) | 10.0, 1.0, 0.1 | 1.0 |
| SVM | class balancing | yes, no | yes |
| RBF kernel | gamma | 1.0, 0.1, 0.01, scale=1 / (n_features * var(X)) | scale |

PCA: Principal Component Analysis

SVM: Support Vector Machine

RBF: Radial Basis Function

**Supplementary Table 3.** Summary table compiled according to DOME (Walsh et al., 2021) recommendations

| **DOME** | Version | 1.0 |
| --- | --- | --- |
| **Data** | Provenance | HUMSAVAR (UniProt Consortium, 2021), accessed on Aug 4th, 2021 and ClinVar (Landrum et al., 2020), accessed on March 29th, 2021. 11,565 protein sequences and 72,429 SRVs total. N_pos_= 30,481 (P/LP SRVs) and N_neg_= 41,948 (B/LB SRVs). Not previously used. |
|  | Dataset splits | N_pos,train_=27,734, N_neg,train_=38,154. N_pos,blind_=2,747, N_neg,blind_=3,794. 42% positives on training set, 42% positives on blind test set. |
|  | Redundancy between data splits | Maximum pairwise sequence identity between training and testing set is 25% on more than 40% alignment coverage. Enforced with MMseqs2 (Steinegger and Söding, 2017) clustering tool. |
|  | Availability of data | Yes, [...] |
| **Optimization** | Algorithm | Support Vector Machines |
|  | Meta-predictions | No |
|  | Data encoding | Variant positions encoded with two protein sequence embeddings, namely ESM-1V (Meier et al., 2021) and ProtTrans T5 XL U50 (Elnaggar et al., 2021). Protein global feature encoding GO terms by means of an ontology embedding model, Anc2Vec (Edera et al., 2022) |
|  | Parameters | p=31,009 |
|  | Features | f=2,400. Feature selection/dimensionality reduction performed by means of a Principal Component Analysis (PCA) of 5,208 original input features. |
|  | Fitting | p is about half N_pos,train_+N_neg,train_=65,888. The risk of over- and under-fitting is very limited. |
|  | Regularization | No. |
|  | Availability of configuration | Yes, hyperparameters in Supplementary Material (Table 2S). |
| **Model** | Interpretability | Black box, as correlation between input and output is masked. |
|  | Output | Binary classification |
|  | Execution time | 75s for a single variation. |
|  | Availability of software | Webserver: [...] |
| **Evaluation** | Evaluation method | 10-fold cross-validation and blind test set. |
|  | Performance measures | Accuracy, Precision, Recall, F1-score, MCC, ROC-AUC. |
|  | Comparison | SIFT (Ng and Henikoff, 2001), PolyPhen-2 (Adzhubei et al., 2010), PROVEAN (Choi et al., 2012), SNPs&GO (Calabrese et al., 2009), MutPred2 (Pejaver et al., 2019) |
|  | Confidence | No |
|  | Availability of evaluation | Yes, [...] |

**Supplementary Table 4.** E-SNPs&GO performances upon ablation of different GO subontologies.

| Input | Q2* | Precision* | Recall* | F1-Score* | ROC-AUC* | MCC* |
| --- | --- | --- | --- | --- | --- | --- |
| Seq | 82.3 | 90.2 | 64.7 | 75.4 | 79.8 | 0.642 |
| Seq + MF | 83.9 | 88.7 | 70.6 | 78.6 | 82.0 | 0.670 |
| Seq + CC | 83.7 | 88.2 | 70.6 | 78.5 | 81.9 | 0.667 |
| Seq + BP | 83.7 | 89.4 | 69.3 | 78.1 | 81.7 | 0.667 |
| Seq + MF + CC | 85.1 | 87.5 | 74.9 | 80.7 | 83.6 | 0.691 |
| Seq + MF + BP | 85.0 | 87.5 | 74.9 | 80.7 | 83.6 | 0.691 |
| Seq + CC + BP | 84.7 | 87.2 | 74.4 | 80.3 | 83.3 | 0.685 |
| Seq + MF + CC + BP | 86.0 | 85.3 | 80.4 | 82.8 | 85.2 | 0.710 |

Seq: ESM-1v + ProtTrans T5

MF: Molecular Function

CC: Cellular Component

BP: Biological Process

*: For scoring index definition, see Section 2.6 of the main paper
